# Assessing the effects of a 3D pathology tissue-processing workflow on downstream molecular analyses

**DOI:** 10.64898/2026.02.12.705570

**Authors:** Elena Baraznenok, Huai-Ching Hsieh, Lydia Lan, Eric Q. Konnick, Sandy Figiel, Srinivasa R. Rao, Dan J. Woodcock, Ian G. Mills, Freddie Hamdy, Jacob E. Valk, Kelly T. Carter, Ming Yu, Thomas G. Paulson, Suzanne Dintzis, William M. Grady, Jonathan T.C. Liu

## Abstract

Non-destructive 3D pathology methods have emerged in recent years with the potential to enhance standard 2D histopathology by greatly increasing the amount of tissue sampled by imaging and by providing volumetric morphological context. Another key advantage is that tissues remain intact, allowing re-embedding after imaging for potential long-term storage and future histological or molecular analyses. However, the impact of 3D pathology protocols on biomolecules — including DNA, RNA, and proteins — and their compatibility with downstream assays, has not been systematically evaluated. Here, we applied a previously optimized 3D pathology protocol — involving deparaffinization, fluorescent H&E-analog staining, optical clearing, and open-top light-sheet microscopy — to formalin-fixed paraffin-embedded (FFPE) specimens of breast, prostate, and head and neck cancer. Following the protocol, tissues were re-embedded in paraffin and compared with paired FFPE controls that did not undergo 3D pathology processing. DNA and RNA were extracted and subjected to quality assessments. Amplifiability was tested by PCR and reverse transcription quantitative PCR (RT-qPCR) of housekeeping genes. Although the results showed a slight decrease in the average yield and increased fragmentation of both DNA and RNA, amplifiability was largely preserved. Sanger sequencing of the PCR products confirmed accurate sequence determinations, while total RNA sequencing indicated that the global transcriptomic profile was largely unchanged. IHC staining of common biomarkers produced comparable signals, suggesting those proteins are well preserved after the 3D pathology workflow. These results demonstrate the feasibility of combining 3D pathology with downstream molecular applications.

## Introduction

Traditional histopathological examinations for disease diagnosis involve cutting fixed tissue into thin two-dimensional (2D) sections mounted on glass slides, where only a small proportion of the specimen is viewed under a microscope without valuable 3D spatial context [1], [2]. By enabling comprehensive sampling and providing volumetric context for thick tissues, non-destructive 3D pathology has emerged as an alternative to conventional 2D pathology and has gained interest in both clinical applications and fundamental research [1], [2]. A number of studies have now demonstrated the value of 3D pathology for improved visual assessments by human observers and for improved diagnostic determinations through diverse computational methods [2], [3], [4], [5], [6], [7], [8]

A non-destructive 3D pathology workflow for formalin-fixed paraffin-embedded (FFPE) specimens has recently been optimized and described. This method, termed Path3D, involves xylene deparaffinization, staining with a small-molecule (rapidly diffusing) fluorescent analog of hematoxylin and eosin (H&E), and optical clearing in ethyl cinnamate to render the tissue optically transparent [9], [10] [11]. After imaging the processed tissues with volumetric microscopy, such as with open-top light sheet (OTLS) microscopy, the resulting two-channel fluorescence datasets may be false-colored (if desired) to mimic the appearance of conventional H&E staining (**Fig. 1a**) [10], [12]. A key advantage of this protocol is that the tissue remains fully intact after 3D imaging, enabling paraffin re-embedding for long-term preservation. Additionally, it allows for potential integration of downstream molecular assays, such as immunohistochemistry or nucleic acid-based analyses, together with 3D image data for future diagnosis or research applications.

**Figure 1.**
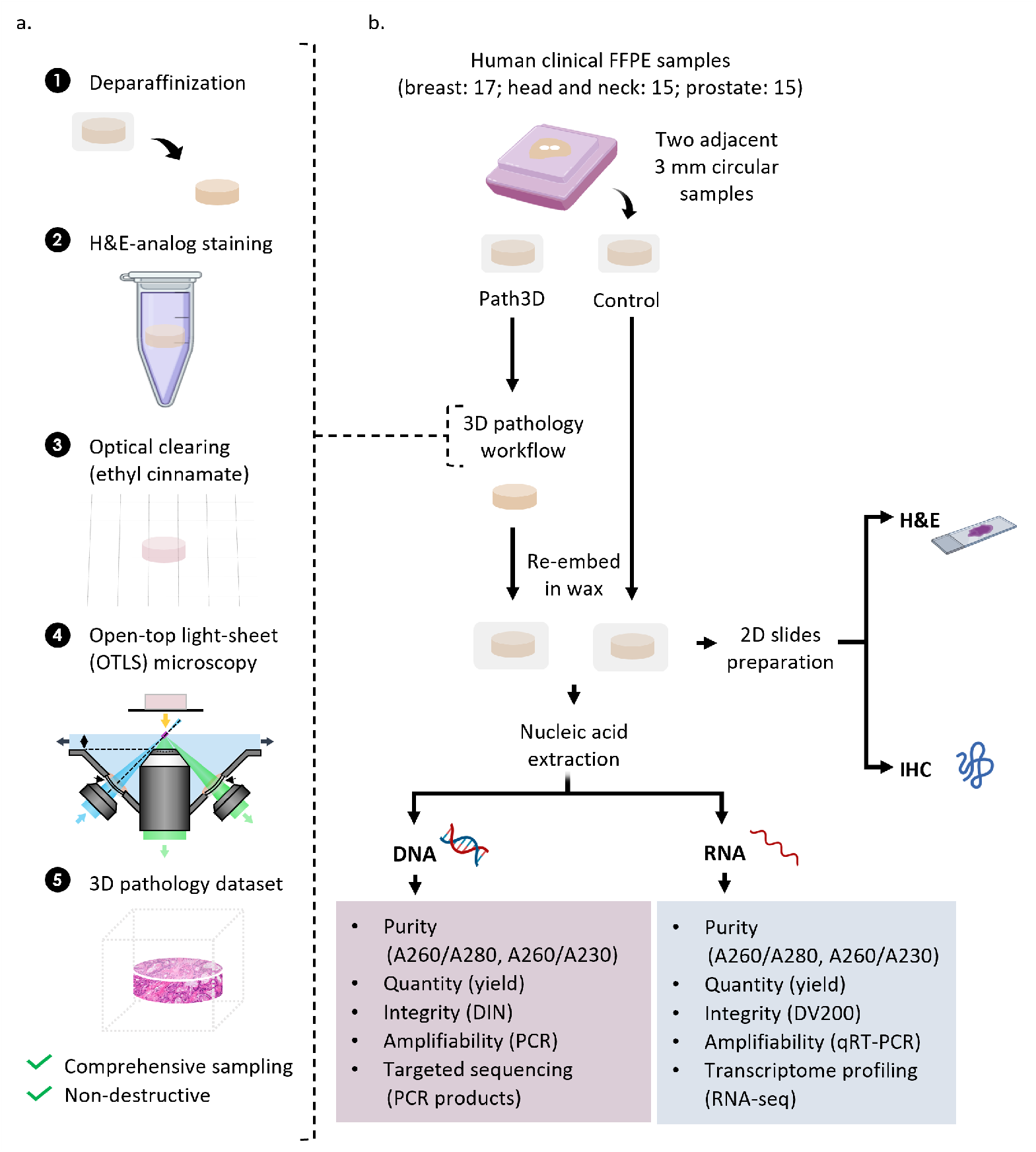
Overview of study design. (a) The 3D pathology (Path3D) workflow used in this study, wherein an FFPE sample undergoes deparaffinization, staining with a fluorescent analog of H&E, optical clearing in ethyl cinnamate, and OTLS microscopy. (b) Schematic of experiments to confirm whether tissues processed by the 3D pathology protocol in (a) are acceptable for downstream molecular assays. Two adjacent circular samples (3 mm in diameter and several mm thick) were punched out from 17 breast cancer, 15 head and neck cancer, and 15 prostate cancer tissue blocks. For each case, one tissue punch sample was designated as the control, while the other was processed using the 3D pathology protocol. Paired punches underwent various molecular assays at the same time (same batch) under identical conditions. Created with Biorender.com

Although Path3D-processed specimens remain physically intact for further analysis, the 3D pathology workflow for large tissue samples involves extended incubation in xylene (for de-paraffinization) and staining buffers containing pH-adjusted ethanol (70%) and sodium chloride. The tissues are also placed in a clearing agent (ethyl cinnamate) for effective 3D imaging. Collectively, these conditions may affect the stability and integrity of biomolecules within the tissue. Assessing the compatibility of the Path3D workflow with downstream molecular assays is of importance, especially for valuable clinical specimens that are archived for various purposes. Here, we evaluated the effects of the 3D pathology tissue processing protocol (Path3D) on DNA, RNA, and proteins across three different tissue types—breast, head and neck, and prostate—by comparing paired processed and unprocessed (control) tissues (**Fig. 1b**). Specifically, we assessed the expression of key protein markers by IHC and the integrity and yield of extracted DNA and RNA. Representative amplification and sequencing assays of DNA and RNA were also performed to confirm their compatibility with standard downstream analyses.

## Methods

### Tissue acquisition

Formalin-fixed, paraffin-embedded (FFPE) tissues were acquired from the Northwest BioSpecimen (NWBioSpecimen) tissue bank at the University of Washington. All processes were conducted in compliance with relevant laws and institutional guidelines and were approved by the University of

Washington Institutional Review Board (IRB No. STUDY00018484, approved on November 8, 2024). The tissue bank collects specimens with patient consent, and all samples were de-identified prior to analysis. Two adjacent circular samples (3 mm in diameter, 1-5 mm tall) were punched out from 17 breast cancer, 15 head and neck cancer, and 15 prostate cancer specimens. For each case, one tissue punch sample was designated as the control, while the other was processed using the 3D pathology protocol.

### 3D pathology tissue processing

One FFPE sample from each pair was deparaffinized by incubation in xylene at room temperature for 48 hours. Following deparaffinization, samples were washed twice in absolute ethanol for 1 hour each and then transferred to 70% ethanol (v/v) for 2 hours. Tissues were stained for 48 hours in 70% ethanol (pH 4) containing 10 mM NaCl, 1:100 eosin (cytoplasm stain; cat. # 3801615, Leica Biosystems) and 1:500 TO-PRO3 (nuclear stain; cat. # T3605, ThermoFisher Scientific). After staining, samples were washed and dehydrated in absolute ethanol twice for 1 hour each, optically cleared in ethyl cinnamate for 2 hours, and finally transferred to fresh ethyl cinnamate for two weeks of storage.

### Paraffin embedding

After completion of the 3D pathology protocol, processed samples were washed twice in absolute ethanol for 1 hour each, followed by a 1-hour wash in 70% ethanol. Control samples were kept in paraffin. Both processed and control tissues were then embedded in new paraffin blocks to enable histological sectioning for downstream analyses. From each FFPE re-embedded block (processed and control samples), ten 10-µm-thick curls were sectioned for nucleic acid extraction. Paraffin embedding and sectioning were performed by the Experimental Histopathology Shared Resources Core at the Fred Hutchinson Cancer Center (FHCC).

### Nucleic acid extraction and quality control (QC)

Nucleic acid extraction and QC measurements were performed by the Biospecimen Processing and Biorepository core at FHCC. DNA and RNA were extracted using the AllPrep DNA/RNA FFPE Kit (Qiagen) according to the manufacturer’s instructions. The initial quantity and purity of both DNA and RNA were assessed with a NanoDrop One spectrophotometer (Thermo Fisher Scientific) based on the 260/280 and 260/230 absorbance ratios.

DNA and RNA concentrations were further quantified using Qubit fluorometric assays. For RNA, aliquots (2-µL) were analyzed with either the Qubit RNA Broad Range Assay Kit or RNA High Sensitivity Assay Kit, depending on the preliminary NanoDrop concentration. For DNA, 2-µL aliquots were measured using the Qubit dsDNA Broad Range Assay Kit or dsDNA High Sensitivity Assay Kit.

DNA and RNA samples with concentrations greater than 7 ng/µL were analyzed for integrity using an Agilent 4200 TapeStation. The DNA Integrity Number (DIN) and DV200 value, suitable for assessment of DNA and RNA integrity from FFPE tissue respectively, were derived from the electropherograms using TapeStation Analysis Software v5.1 (Agilent Technologies).

### DNA/RNA amplification tests

Following the initial QC of the nucleic acids, DNA and RNA amplification tests were performed at the Genomics core at FHCC. The DNA amplifiability of the samples was tested using a PCR assay for two housekeeping genes, *COL2A1* (primer: Hs00570192_CE, Thermo Fisher Scientific) and *GAPDH* (primer: Hs00828901_CE, Thermo Fisher Scientific). Aliquots of 15 ng DNA were used, with two technical replicates per sample. Amplification was accomplished using the DreamTaq™ Hot Start PCR Master Mix (Thermo Fisher Scientific), according to the manufacturer’s instructions, on an ABI QuantStudio5 Real Time PCR System. DNA PCR products were purified using Solid Phase Reversible Immobilization (SPRI) magnetic beads, quantified with Qubit, and subsequently analyzed using the D1000 DNA ScreenTape assay. Electropherograms and housekeeping gene product concentrations were analyzed using TapeStation Analysis Software v5.1 (Agilent Technologies).

RNA amplifiability was tested using a reverse transcriptase (RT) qPCR assay for *GAPDH*. A forward primer and two reverse primers, previously reported in the literature [13], were used to amplify different amplicon sizes. The sequences were as follows: forward—5′-CCACATCGCTCAGACACCAT-3′; 71 bp—5′-GTAAACCATGTAGTTGAGGTC-3′; 153 bp—5′-ACCAGGCGCCCAATACG-3′. RT-qPCR was performed using SuperScript™ III Platinum™ SYBR™ Green One-Step Kit (Thermo Fisher Scientific), following the manufacturer’s protocol, with 50 ng RNA per reaction and two technical replicates per biological sample. The cycle threshold (Ct) values and melt curves were analyzed and compared between paired samples.

### Targeted sequencing of DNA

The *GAPDH* and *COL2A1* DNA PCR products from the amplification tests with Qubit concentration greater than 0.5 ng/uL were sequenced by the Genomics Core at FHCC. To prepare the reaction, 5 µL of the PCR product (diluted to 0.5-5 ng/µL) was combined with 3 uL of the corresponding forward primer (*COL2A1* or *GAPDH*). Sequencing and fragment analysis was performed on an Applied Biosystems (ABI) 3730xl DNA Analyzer. Poor-quality reads at the ends of the raw sequence data were trimmed in python using a quality score threshold of 30 for a sliding window size of 10 nucleotides [14]. From the trimmed sequences, the percentage of bases with quality over 30 (Q30) was calculated, along with the average quality score, the length of the sequence, and the pairwise alignment similarity percentage to a reference genome (GRCh38).

### Total RNA sequencing and analysis

RNA from two pairs per tissue type was sequenced at the FHCC Genomics core. Whole transcriptome libraries were prepared using the Illumina Stranded Total RNA Prep Kit and sequenced on the Illumina NovaSeq X Plus platform in paired-end 150 bp mode, generating approximately 90–107 million reads per sample. Raw FASTQ files were quality-checked using FastQC (v0.12.1) [15], trimmed with Cutadapt (v4.9) [16], and aligned and quantified using STAR (v2.7.10b) [17] against the human reference genome (GRCh38). FastQC was also used to evaluate sequence quality after trimming. Gene body coverage analysis was performed using RSeQC (v5.0.4) [18]. Gene-level raw counts were normalized using variance stabilizing transformation (VST), and pairwise global transcriptomic correlations were calculated using Spearman’s rank correlation in the DESeq2 R (v4.5.2) package (v1.50.0) [19].

### H&E and immunohistochemistry

Three pairs of samples per tissue group (breast, head and neck, prostate) underwent histological examination to qualitatively assess H&E and IHC. One H&E slide was prepared per sample. From each of the selected breast samples, one slide was stained with an antibody for estrogen receptor (ER) (1:100, cat. # ab16660, abcam), and one slide was stained for HER2/neu (1:100, cat. # ab16662, abcam). The head and neck samples were stained for pan-cytokeratin (1:300, cat. # ab7753, abcam), while the prostate samples were stained for NKX3.1 (1:250, cat. # 35-9700, Thermo Fisher Scientific).

All H&E and IHC work was conducted at the Histology and Imaging Core (HIC) in the Department of Comparative Medicine at the University of Washington. For IHC, slides were processed on the Leica BOND RX Fully Automated Research Stainer (Leica Biosystems). Following deparaffinization, antigen retrieval was performed in either EDTA (breast: 10 min, H&N: 20 min) or citrate (prostate: 10 min) at 100°C. The BOND Polymer Refine Detection system reagents (Leica Biosystems) were then applied sequentially per the manufacturer’s directions to complete the antibody stain, DAB detection and hematoxylin counterstain. Each slide was scanned at 20X to produce a whole-slide image.

### Statistics

Paired t-tests were used to determine the statistical significance of differences in the means between the 3D pathology processed and control groups for quantitative nucleic acid metrics. A p-value <0.05 was considered statistically significant. Box plots were generated to depict the arithmetic mean, interquartile range, and standard deviation.

## Results

### Impact of Path3D protocol on nucleic acid extraction

Approximately one month after the 3D pathology process, DNA and RNA were successfully extracted from all FFPE tissue pairs (15 head and neck, 15 prostate, and 17 breast cancer specimens). For DNA, the average A260/A280 ratios were above 1.80 in both the processed and control groups, suggesting low protein contamination (**Fig. 2a**). The A260/A230 ratios showed greater inter-sample variation but still averaged above 1.70 and were similar between groups, indicating acceptable purity (**Fig. 2b**). For RNA, the mean A260/A280 ratios in both groups also exceeded 1.80 and remained comparable (**Fig. 2d**). In contrast, the A260/A230 ratios were lower in the Path3D tissues (1.38 vs control 1.52), likely due to residual organic compounds introduced during the 3D pathology protocol (**Fig. 2e**).

**Figure 2.**
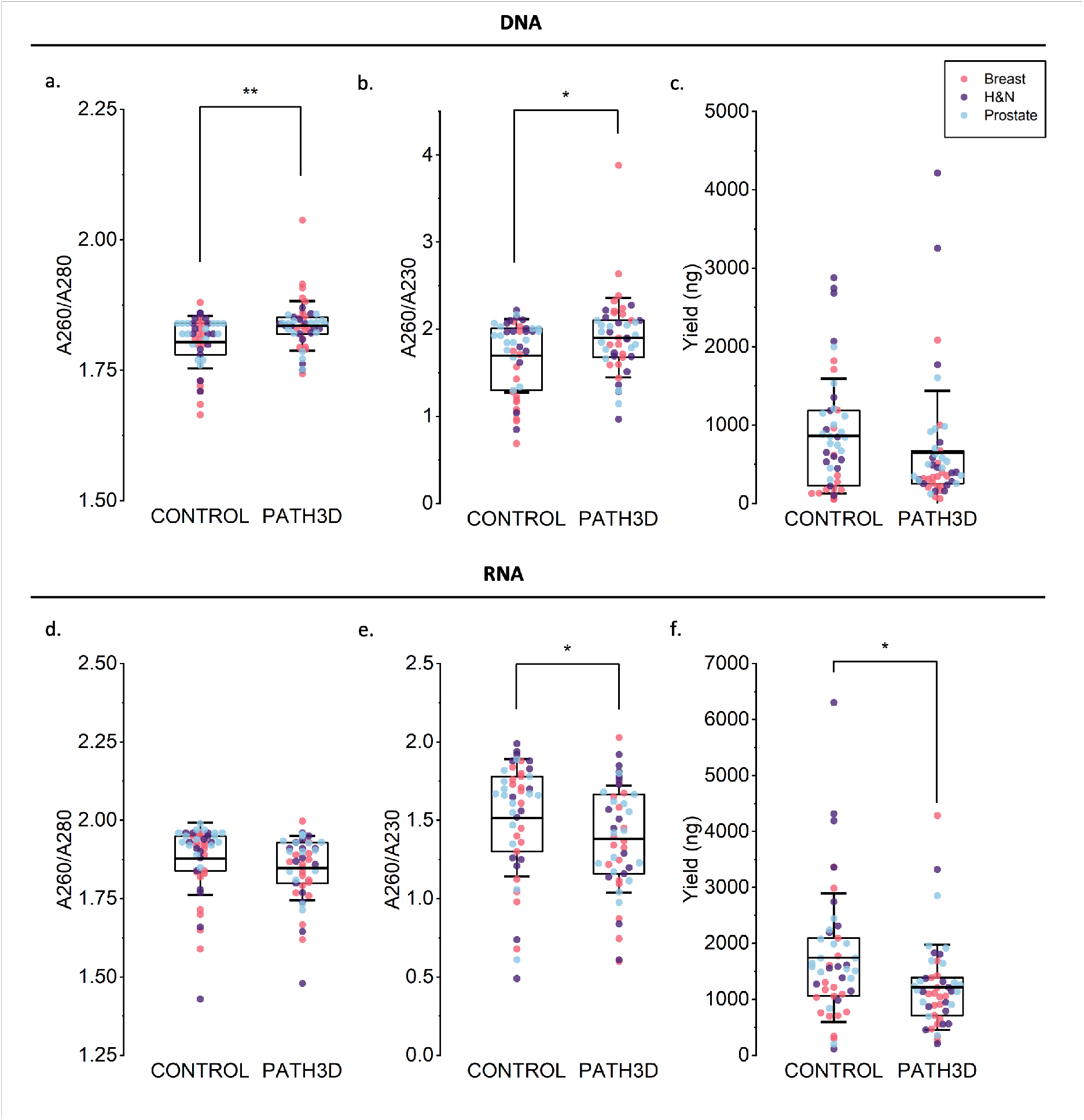
Distributions of DNA and RNA purity, quantity and integrity results. Points show individual breast (red), prostate (blue), and head and neck (dark purple) cancer samples. (a) DNA A260/A280 absorbance ratio. (b) DNA A260/A230 absorbance ratio. (c) Quantity of DNA, measured by Qubit. (d) RNA A260/A280 absorbance ratio. (e) RNA A260/A230 absorbance ratio. (f) Quantity of RNA, measured by Qubit. ^*^ p<0.05, ^**^ p<0.001

Despite considerable inter-sample variability, nucleic-acid yields were comparable across tissue types and were therefore analyzed collectively. On average, the processed samples yielded less DNA (649 ng vs. 860 ng) and RNA (1214 ng vs. 1741 ng) (**Fig. 2c, f**), suggesting gradual degradation of nucleic acids during the workflow. Average DNA and RNA quantities varied depending on tissue type, with breast cancer specimens yielding the least nucleic acid, but the relative amount extracted from Path3D compared to control was always lower (per tissue type, control vs. Path3D: breast, 1790 ng vs. 1567 ng; head and neck, 3530 ng vs. 2087 ng; prostate, 2593 ng vs. 1977 ng). Overall, the 3D pathology protocol modestly reduced RNA purity (<10% decrease) and DNA/RNA recovery (24% and 30% decreases, respectively), while extracted material remained sufficient for downstream molecular analyses.

### Impact of Path3D protocol on DNA and RNA integrity

Next, integrity scores for both DNA and RNA were analyzed to evaluate the quality of extracted nucleic acids. For DNA, the average DIN scores were 3.2 for the Path3D group and 4.4 for the control group, suggesting a statistically significant increase in DNA fragmentation during processing (**Fig 3a**). This was further supported by the representative gel images of paired samples, showing a shift of DNA peaks toward shorter fragment lengths in the Path3D group for most samples (**Fig. 3b**). For RNA, RNA integrity number (RIN) scores were low across all samples (RIN <2.5) because archived FFPE samples always contain highly fragmented RNA. Hence, the DV200 percentage—the percentage of RNA fragments longer than 200 bp—was used to compare the two groups. On average, the DV200 percentages were reduced in the Path3D processed samples (50.84%) compared to the control group (58.54%), suggesting increased RNA fragmentation with 3D pathology processing (**Fig. 3c**). Consistent with the DNA results, RNA gel lanes exhibited smeared and downward-shifted signals, indicating increased fragmentation (**Fig. 3d**). Collectively, these results suggest that DNA and RNA integrity was already compromised in the FFPE samples and further decreased following 3D pathology processing.

**Figure 3.**
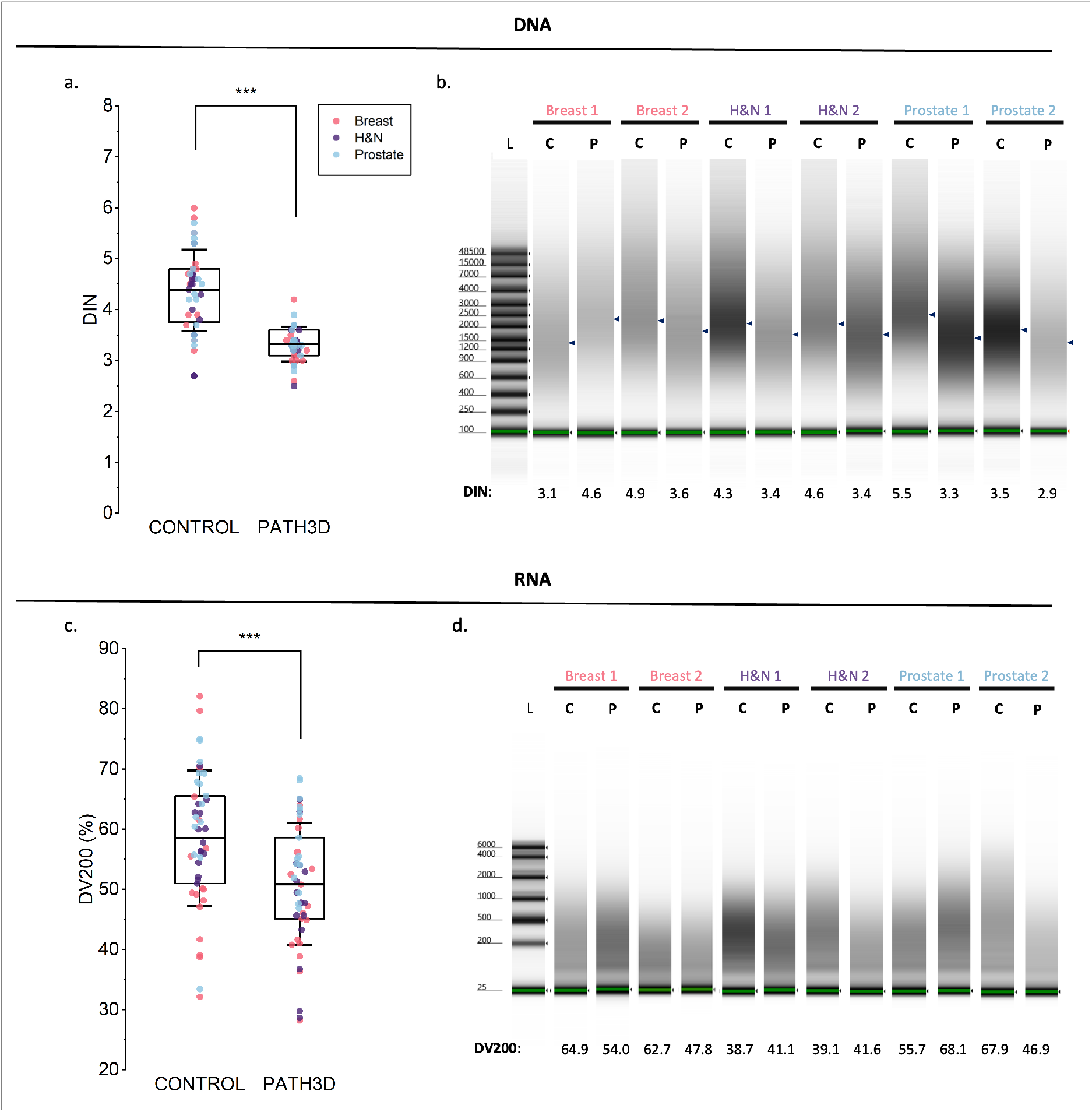
Nucleic acid integrity results for breast (red), prostate (blue), and head and neck cancer specimens (green). L = ladder, C = control, P = Path3D processed. (a) Distribution of DIN Scores. (b) TapeStation gel images for DNA. Peak sample concentration is denoted by marker. (c) Distribution of DV200 percentages (d) TapeStation gel images for RNA. ^***^ p<0.0001

### Successful DNA amplification and sequencing following Path3D processing

After confirming modest degradation and fragmentation, we next examined whether the extracted nucleic acids remained compatible with common downstream molecular assays. PCR amplification was performed for two housekeeping genes, *GAPDH* (228 bp) and *COL2A1* (274 bp). Representative TapeStation gel electrophoresis images showed clean bands for both groups with little to no off-target signals, indicating specific amplification (**Fig. 4e,f**). The average product lengths matched the expected amplicon sizes and did not significantly differ between groups for both *GAPDH* (232 bp for both) and *COL2A1* (274 bp vs. 275 bp). Quantitatively, the average TapeStation peak concentrations for the *GAPDH* amplicon were 2.34 ng/µL and 2.37 ng/µL for the Path3D and control groups (**Fig. 4a**), while those for the *COL2A1* amplicon were 3.34 ng/µL and 4.54 ng/µL (**Fig. 4g**).

**Figure 4.**
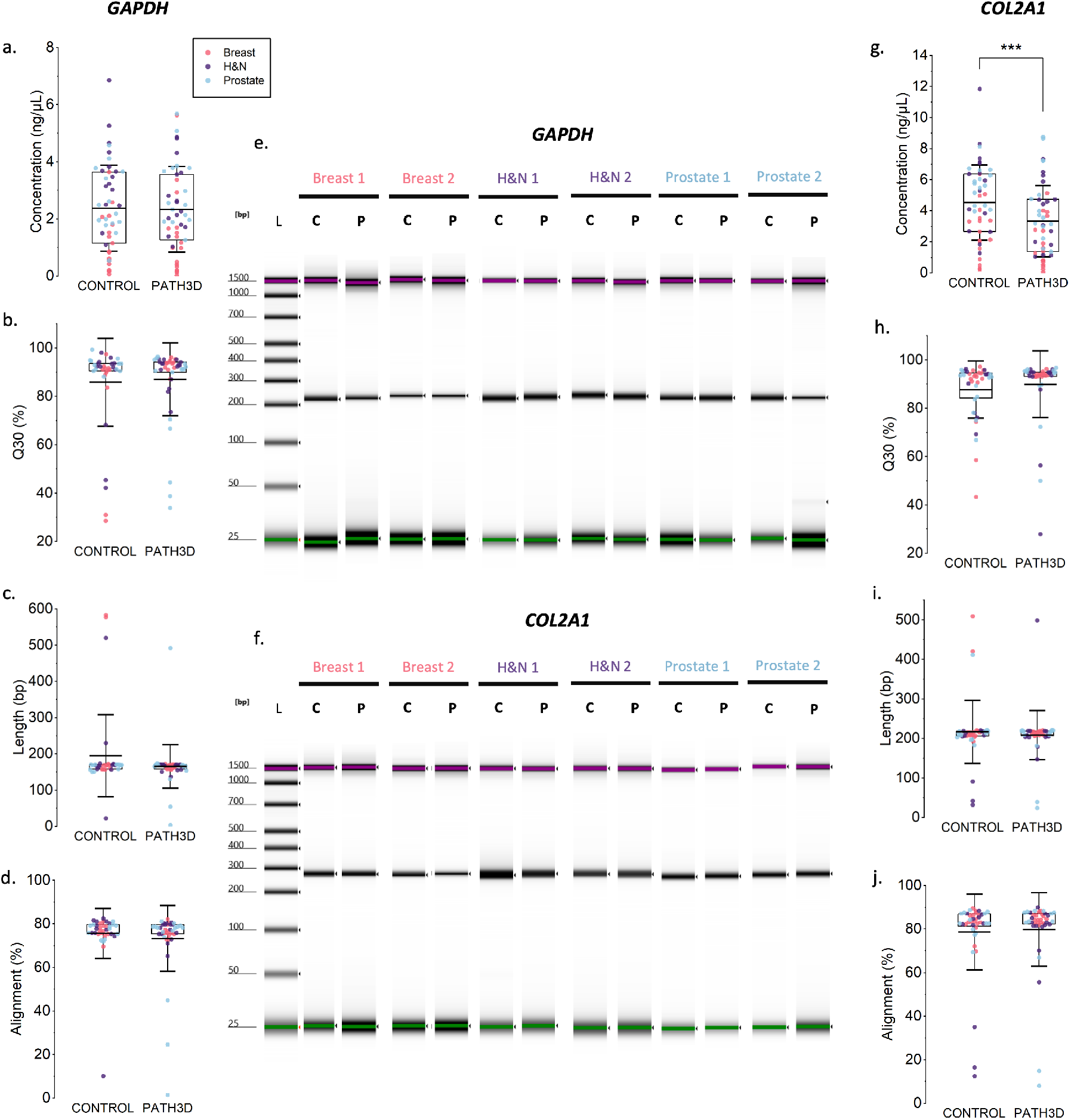
DNA amplification and sequencing test results. L = ladder, C = control, P = Path3D processed. PCR tests were performed on sample pairs in which at least one replicate had a product concentration ≥ 0.5 ng/µL—a threshold met by most samples. (a) Distribution of *GAPDH* PCR target product concentration (n=47). (b) Distribution of Q30 values for *GAPDH* PCR product sequencing (control: n=37, Path3D: n=42). (c) Distribution of sequence length of *GAPDH* PCR products (control: n=37, Path3D: n=42). (d) Distribution of similarity of *GAPDH* product to reference sequence (control: n=37, Path3D: n=42). (e) Screentape gel image for *GAPDH* PCR products. (f) Screentape gel image for *COL2A1* PCR products. (g) Distribution of *COL2A1* PCR target product concentration (n=47). (h) Distribution of Q30 for *COL2A1* PCR product sequencing (control: n=40, Path3D: n=41). (i) Distribution of sequence length of *COL2A1* PCR products (control: n=40, Path3D: n=41). (j) Distribution of similarity of *COL2A1* product to reference sequence (control: n=40, Path3D: n=41). ^***^ p<0.0001

DNA usability was further assessed using Sanger sequencing of PCR products. As FFPE-derived DNA can have variable quality, a successful sequence was defined as one that could be trimmed by the sequence quality–based algorithm (described in the Methods section), indicating sufficient read length and base quality. The number of successful sequences was comparable between the two groups for both *GAPDH* (control: n=37, Path3D: n=42) and COL2A1 (control: n=40, Path3D: n=41), and these sequences were further used for analysis of sequencing metrics. For both genes, the average Q30 values were uniformly high and comparable between the Path3D and control groups—87% vs. 86% for *GAPDH* (**Fig. 4b**) and 90% vs. 88% for *COL2A1* (**Fig. 4h**)—indicating that 3D pathology processing did not affect sequencing quality. The average sequence lengths and alignment similarity to the reference sequences were also largely comparable for *GAPDH* (control: 195 bp, 76% similarity; Path3D: 165 bp, 73% similarity) and *COL2A1* (control: 217 bp, 78% similarity; Path3D: 208 bp, 80% similarity) (**Fig. 4c,d,i,j**). Taken together, these results indicate that 3D pathology processing did not substantially affect DNA amplification and sequencing, although a lower yield was observed for *COL2A1*.

### Retention of RT-qPCR amplifiability within FFPE-compatible amplicon sizes following Path3D processing

RNA amplifiability was assessed using RT-qPCR assays targeting two *GAPDH* regions, 71 bp and 153 bp, representing short and moderately longer FFPE-compatible targets. While the average Ct value for both amplicons was slightly higher in the Path3D-processed group compared with the control, corresponding to approximately 22% and 28% lower amplification in 71 bp and 153 bp amplicons, respectively, both amplicons remained robustly detectable (**Fig. 5a**). These results indicate preservation of RT-qPCR amplifiability within the size range relevant for FFPE-derived RNA and motivated subsequent evaluation of global transcriptome preservation by RNA sequencing.

**Figure 5.**
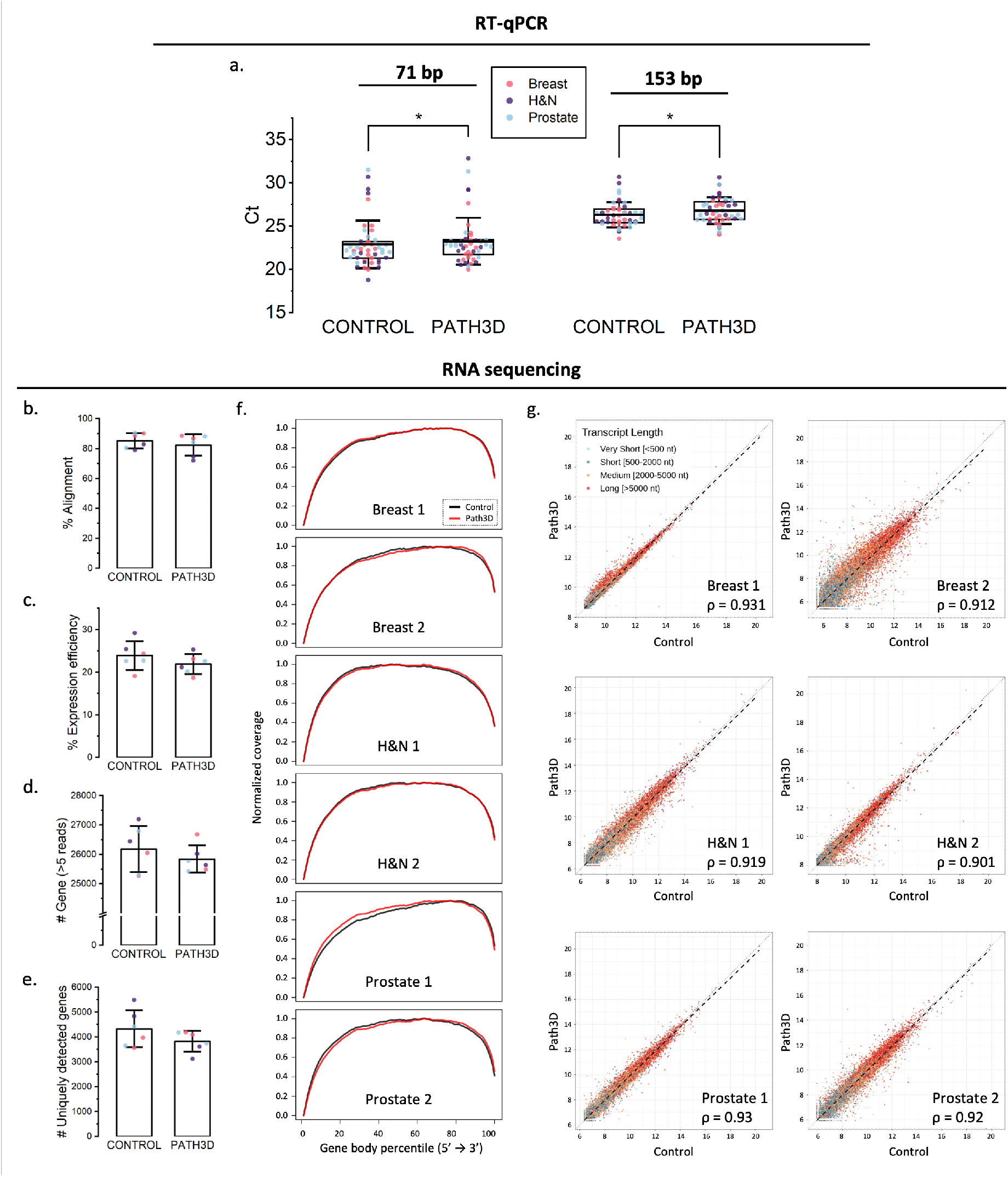
RNA amplification and sequencing test results. (a) Average Ct value for amplicons of length 71 bp and 153 bp. *p<0.05 (b) Average percent of sequence reads aligned to reference genome. (c) Average percent expression efficiency (exonic reads). (d) Average percent of gene counts of genes with more than 5 reads. (e) Number of uniquely detected genes in only the control group or only the Path3D group. (f) Pairwise normalized gene body coverage profiles. (g) Scatterplots showing the relationships between control (x) and Path3D (y) of 6 specimens, colored by transcript length, with Spearman correlation coefficients (ρ). ^*^p<0.05, ^**^p<0.001

To evaluate whether transcriptomic profiles were globally preserved, we performed total RNA sequencing. Six pairs of samples with high DV200 (control average: 67.1%, Path3D average: 59.8%) were sequenced. As a result, the average percentage of RNA sequencing reads successfully mapped to the human reference genome was high in both the control (85.3%) and Path3D (82.4%) groups (**Fig. 5b**), suggesting that library preparation captured transcripts broadly in both groups, with successful alignment. Transcript coverage of aligned reads was further evaluated using gene body coverage analysis, which represents average normalized coverage along transcript bodies. Across the six pairs, most Path3D samples showed mildly right-shifted coverage curves relative to matched controls, with largely preserved transcript coverage profiles (**Fig. 5f**). In addition, the average percentages of those reads mapping to exonic regions (expression efficiency) were 23.9% and 21.9% (**Fig. 5c**) and the average total gene counts were 26,179 and 25,843 (**Fig. 5d**) for control and Path3D, respectively. Although the integrity of input RNA was slightly lower in the Path3D group, the differences in these three RNA-sequencing QC metrics were modest, indicating minimal impact on overall RNA-sequencing performance.

To further characterize transcriptomic similarity between Path3D and control samples, pairwise analyses were performed. The average numbers of genes detected exclusively in the control group and exclusively in the Path3D group were 4,321 and 3,822, respectively, indicating similar gene-dropout counts and preservation of low-abundance transcripts following Path3D processing (**Fig. 5e**). Pairwise VST-based Spearman correlation analysis revealed high similarity in global gene-expression profiles for Path3D and matched control samples from the same tissue blocks (ρ > 0.9 for all six pairs), with minor differences likely reflecting tissue heterogeneity (**Fig. 5g**). The VST-based scatter plots showed a symmetric distribution along the identity line, including low-abundance transcripts, indicating that no systematic loss or compression of weak signals was attributable to Path3D processing. Additionally, no color separation orthogonal to the identity line was observed (with points color-coded by transcript length), suggesting no biased expression-level shift toward either shorter or longer transcripts. Overall, these results suggest that RNA following Path3D processing remains suitable for RNA sequencing.

### Preservation of tissue morphology and protein markers in H&E and IHC assays

H&E and IHC staining were performed to assess tissue morphology and protein marker preservation following the 3D pathology workflow. **Figure 6** presents representative H&E comparisons between control and Path3D samples for two pairs per tissue type. The slight shrinkage and marginally darker staining in the Path3D samples, possibly due to increased tissue density from shrinkage, were within the interpretable range based on pathologists’ review. A comparison of ER and HER2 IHC for breast samples (**Fig. 6a**), pan-cytokeratin for head and neck samples (**Fig. 6b**), and NKX3.1 for prostate samples (**Fig. 6c**) is also shown. No residual H&E-analog staining from the 3D pathology workflow was observed in the IHC images. IHC staining intensity was largely comparable between control and Path3D samples, and the stain matched the intended target. Collectively, these findings demonstrate that 3D pathology processing preserves tissue architecture and antigen detectability, supporting compatibility with standard histopathologic and immunohistochemical analyses.

**Figure 6.**
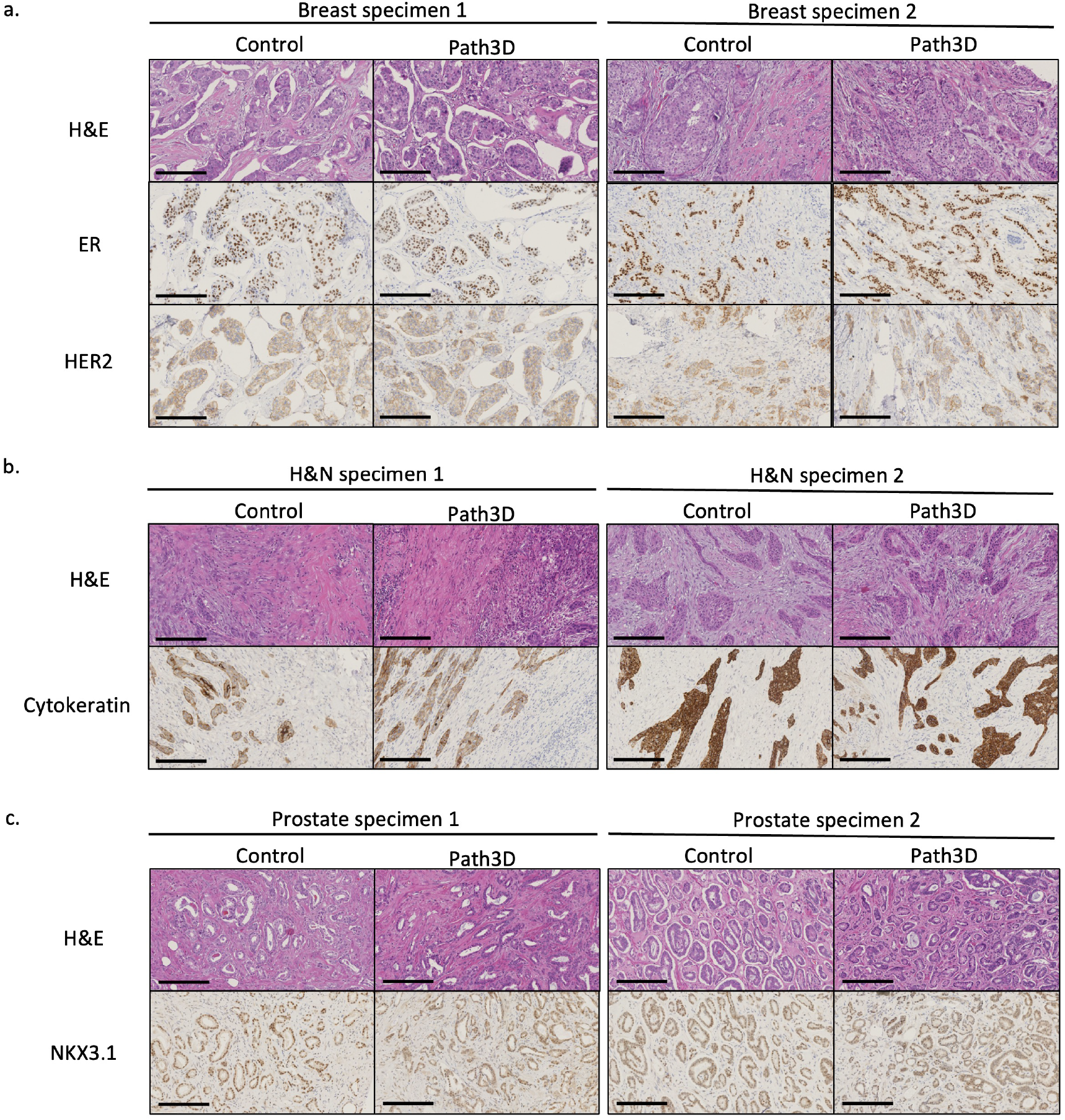
H&E and IHC image comparisons, 20X for (a) breast specimens: diffusely positive estrogen receptor staining for invasive ductal carcinoma (specimens 1 and 2); faint, incomplete (specimen 1) and weak to moderate variably complete (specimen 2) membrane staining for Her2 protein, (b) head and neck specimens: moderate membrane (specimen 1) and strong membrane (specimen 2) staining for pan-cytokeratin in invasive squamous cell carcinoma, and (c) prostate specimens: positive moderate-to-strong intensity staining for Gleason pattern 3 (specimen 1) and Gleason pattern 3+4 (specimen 2). Scale bar: 200 µm.

## Discussion

Non-destructive 3D pathology offers several compelling advantages over standard 2D pathology, including much greater sampling of specimens, comprehensive 3D spatial context, and a slide-free (sectioning-free) process that keeps tissues intact for downstream uses [20], [21]. Here, we further evaluate the effects of the protocol on biomolecular integrity. While nucleic acid yields and integrity in the 3D pathology processed samples were slightly affected in comparison to corresponding unprocessed FFPE control samples, compatibility with downstream molecular and histological assays — including PCR, Sanger sequencing, qPCR assays, total RNA sequencing, and IHC staining — were largely preserved.

The observed reduction in nucleic acid recovery and integrity is not unexpected, given the extended processing periods (∼1 week for Path3D plus ∼2 weeks for molecular analysis) during which samples were subjected to various staining, clearing, and buffer-exchange steps. Such prolonged processing can lead to enzymatic degradation, chemical modification, or partial loss of nucleic acids [22], [23]. It should be noted that not all buffers and equipment were maintained under nuclease-free or proteinase-free conditions in this study, and that incubation times were quite long in certain buffers (e.g. there was ∼1 month of time between Path3D completion and the completion of nucleic acid extraction). Therefore, additional optimization should further improve biomolecular preservation in the 3D pathology workflow. For example, prior studies have shown that certain additives and buffer conditions can reduce DNase and RNase activities, and that minimizing processing/incubation times can also help to maintain nucleic acid integrity [24].

A limitation of this study is that both the PCR and IHC analyses focused on commonly expressed genes and protein markers. For DNA, assessing additional genomic targets could provide further insight into how the protocol influences genomic DNA integrity. The observed variation in amplification efficiency between the two housekeeping genes, *COL2A1* and *GAPDH*, suggests that degradation and fragmentation introduced by the protocol may affect certain genomic regions differently, depending on their location or structural characteristics. For IHC, future studies may benefit from expanding to a broader antibody panel or incorporating spatial proteomic techniques to better evaluate the impact on a wider range of proteins and to assess compatibility with standard spatial omics methods.

In summary, our findings support the feasibility of integrating a non-destructive 3D pathology protocol into molecular workflows without significantly compromising molecular-assay performance. To our knowledge, this represents the first comprehensive and quantitative evaluation of a 3D tissue processing workflow for clinical specimens in terms of compatibility with downstream molecular assays. Given the growing adoption of optical clearing across biomedical research, our results also provide insights into the potential impact of ethyl cinnamate–based clearing, one of the most widely used clearing methods [11], [25], on different biomolecules. As molecular data becomes increasingly important in diagnostic pathology, protocols that preserve both tissue architecture and nucleic acid quality are essential. As mentioned, further steps to optimize the Path3D protocol, such as using DNase and RNase-free reagents and minimizing incubation times in certain reagents, are likely to further reduce biomolecule degradation. Overall, this work highlights the potential of 3D pathology workflows to enable comprehensive imaging of intact tissues while retaining valuable samples for long-term storage and subsequent molecular and histological analyses.

## Ethics declarations

### Competing interests

J.T.C.L. is a cofounder, equity holder, and board member of Alpenglow Biosciences, Inc., which has licensed the 3D pathology technologies developed in his lab, including patents related to open-top light-sheet (OTLS) microscopy. W.M.G. is a consultant for Guardant Health, Karius, and DiaCarta, and receives research support from Lucid Diagnostics.

## Acknowledgements

This work was supported by funding from the National Institutes of Health (NIH) through R01CA268207 (J.T.C.L.), R01DK138948 (J.T.C.L., W.M.G., T.G.P), and U54DK137328 (J.T.C.L.). NIH also supported through U01AG077920 (W.M.G., M.Y.), U2CCA271902 (W.M.G., M.Y., T.G.P.), U54CA274374 (W.M.G., M.Y.), R21CA259687 (T.G.P.), R01CA266386(T.G.P.), R01CA140657 (T.G.P.), R01CA270235 (T.G.P.), and R50CA233042 (M.Y.). M.Y. is also supported by the Kuni Foundation. Funding for W.M.G. is also provided by the Prevent Cancer Foundation, Cottrell Family Fund, and Listwin Foundation. Support was also provided by the Prostate Cancer United Kingdom (PCUK) charity through MA-ETNA19-005 (F.C.H., I.G.M, S.F., S.R.R., D.J.W., J.T.C.L.). The John Black Charitable Foundation also supports the work of I.G.M. S.F. and I.G.M. are in addition funded by a Wellcome Trust Bioimaging Award (313477/Z/24/Z). Additional support was provided by the Department of Defense (DoD) Prostate Cancer Research Program: W81XWH-20-1-0851 (J.T.C.L.), W81XWH-18-10358 (J.T.C.L.), and the Advanced Research Projects Agency for Health (ARPA-H) D24AC00357 (J.T.C.L.).

